# Why budding yeast overrides the DNA damage checkpoint

**DOI:** 10.1101/2025.07.30.667764

**Authors:** Roxane Oesterle, Sahand Jamal Rahi

## Abstract

Checkpoints arrest biological processes and enhance chances for error correction. In many species ranging from budding yeast to human, checkpoints are eventually overridden despite persistent damage. Whether checkpoint override serves a biological function remains unclear. Here, we investigate this question in the context of the DNA damage checkpoint (DDC) in budding yeast. To demonstrate that DDC override increases fitness, we pursued a novel approach: To avoid inherent ambiguities when comparing genetic mutants, we instead employed a light-controlled trigger to finely tune the timing of checkpoint override in a consistent wild-type DDC and DNA repair background. We show that override is beneficial and wild-type override timing maximizes fitness. We formulate two specific hypotheses to explain the fitness benefit: i) override enables multiple rounds of replication, including of broken chromosomal fragments, statistically increasing the chance of at least one successful repair; or ii) override may enhance specific DNA repair pathways. Testing the first hypothesis, we tracked broken chromosome fragments using an optogenetic reporter and found their segregation pattern to be inconsistent with a probabilistic increase in post-override repair opportunities. To test the second hypothesis, we dynamically depleted key repair pathway proteins – individually and combinatorially – without interfering with the establishment of checkpoint arrest. Strikingly, we found that proteins involved in microhomology-mediated end joining (MMEJ) substantially enhanced override-associated break repair. Together, these results provide direct evidence of a fitness advantage conferred by checkpoint override and uncover MMEJ-associated repair proteins as the mechanistic basis.

## Introduction

Cell cycle and developmental checkpoints detect errors and arrest cellular processes, which provides an opportunity for correction and promotes genomic and organismal integrity. Yet, across diverse species and contexts, biological systems have been observed to override checkpoints in the continued presence of errors. Examples include the DNA damage checkpoint (DDC), which has been reported to be overridden (or ‘leaky’) in the presence of persistent DNA damage in *Saccharomyces cerevisiae* ^1,2^, *Caenorhabditis elegans* embryos^3^, *Xenopus* egg extract^4^, and both cancerous^5,6^ and non-cancerous^7–10^ human cells. Similarly, prolonged spindle attachment defects can lead to “slip-page” through the spindle assembly checkpoint in both yeast and human cells^11–16^. In insects, developmental checkpoints have been reported to delay – but not abort – transitions under sustained tissue damage^17,18^.

Despite evolutionary conservation and physiological importance of checkpoints, the biological function of checkpoint override remains unresolved. Even for the more extensively studied case of DDC override in budding yeast^19^, different perspectives persist about whether override is an adaptive, fitness-enhancing response^20^; detrimental to genome integrity^21^, potentially due to mechanical or signaling failure after prolonged arrest; or of no consequence to organismal fitness^22^. A ‘null hypothesis’ posits that prolonged arrest leads to abnormally enlarged cells and thereby causes checkpoint regulators to become too diluted, weakening checkpoint signaling. Other components of the cell could likewise deteriorate under arrest, leading to checkpoint failure. The fact that few cells or animals survive after override further argues against any adaptive value.

A rigorous test of the biological function of override requires a quantitative comparison of fitness between organisms with wild-type and altered override behavior. However, designing and executing such a test presents conceptual and technical challenges: From a conceptual standpoint, fitness is a complex organismal attribute that manifests in a multi-faceted ecological context over evolutionary timescales. Vastly different aspects of organismal behavior influence fitness and may in principle need to be evaluated. Methodologically, most studies of override rely on constitutive mutations or molecular perturbations, e.g., *cdc5-ad* in budding yeast, that block the process. Yet, in complex systems such as checkpoints and repair pathways, these interventions can have pleiotropic effects. Distinguishing the direct impact of override disruption from secondary consequences is extremely challenging, especially when the perturbation is permanent. One can overcome these limitations by avoiding comparisons between distinct genetic backgrounds altogether and, instead, manipulating override timing in a controlled, reversible manner within the same background. Such a strategy allows more precise attribution of fitness effects to override itself. Finally, the case for a functional role of override would be buttressed by showcasing a mechanism by which this benefit arises.

Thus far, the data do not permit a definitive conclusion^19^. Indirect evidence in favor of a biological function is provided by the existence of mutations that abrogate DDC override in budding yeast^2,23–25^, suggesting that the phenomenon may have been under positive selection. However, in direct tests, adaptation-competent (*CDC5*) and -incompetent (*cdc5-ad*) haploid cells showed no difference in sensitivity to X-ray radiation^26^. In diploids, high doses of radiation or knock-out of homologous recombination (*rad52*Δ) were needed to observe a difference since individual chromosomes can be dispensable. We recently showed that the timing of DDC override in the presence of one or multiple DSBs agrees well with theoretically optimal override times, which maximize the expected number of offspring by balancing the chances of DNA break repair and of survival^20^. Other modeling work has shown that heterogeneity in override times should provide a benefit in fluctuating environments^27^. On the other hand, DDC override is also associated with genomic instability^21,26,28^, which is usually considered to be detrimental^26^ but leads to resistance to genotoxic stress in a repair-deficient background^28^.

In addition to the need to settle whether there is a benefit to checkpoint override, the underlying reasons need to be explored. While override could affect fitness in ways that would be extremely challenging to prove experimentally, e.g., by increasing genetic instability and thereby chances for beneficial evolutionary innovations^27^, we primarily investigate the impact on DNA break repair, which is necessary for survival. To formulate testable mechanistic hypotheses for a potential benefit of DDC override, we analyzed the difference between cells with double-strand DNA breaks (DSBs) pre- and post-override. Two differences stand out: Since override is often the first of a small number of cell cycles (all post-override progeny eventually die unless at least one cell repairs the DSBs), there are more copies of the genome, distributed into more cells, after override. Another difference is that an undivided mother-daughter pair is arrested pre-anaphase before override, while after override the budded cell completes mitosis and both mother and daughter cells can in principle go through other cell cycle stages, including G1, S phase, and mitosis. These considerations lead to two distinct hypotheses:

1. Statistical benefit hypothesis: The multiple new copies of the genome that segregate into different cells could in principle increase the probability that in at least one of the cells, the DSB is fixed, leading to a greater chance of survival.
2. Repair benefit hypothesis: In the post-override cell cycle stages, different repair pathways could become accessible, increasing the chance of repair and survival.

Repair of the DSBs is critical for survival and proceeds by four main pathways^29^: non-homologous end-joining (NHEJ), homologous recombination (HR), single-strand annealing (SSA), or alternative end joining (alt-EJ). In NHEJ, minimally processed break ends are ligated together. NHEJ predominantly takes place in G1 phase and requires the binding of the Ku70-Ku80 dimer to the broken ends, preventing end resection and recruiting DNA ligase 4 complex subunits Dnl4 and Nej1. The three other pathways involve end resection. HR relies on a homologous sequence as a template for repair and mainly occurs in S/G2. It is dependent on Rad51 for strand invasion and Rad52 for loading of Rad51 on DNA. SSA repairs the breaks by annealing repeated homologous sequences from either side of the break using Rad52. Microhomology-mediated end joining (MMEJ) is plausibly the best studied type of alt-EJ^30,31^. Similar to SSA, MMEJ aligns DNA break ends but uses microhomologies (5-25 base pairs^32^) and is independent of Rad52. SSA and MMEJ both also require Rad1-Rad10. They preferentially occur in S/G2 phase or mitosis and usually result in deletions^29,30,33,34^. We will leverage the different genetic requirements for repair pathways^35^ (**Table 1**) in order to disable them differentially and dynamically. In this manner, we can explore the repair benefit hypothesis by assessing the contribution of each pathway to override-dependent survival benefits, if any.

**Table 1:** Repair proteins required (+), moderately important (x), or not required (–) for each repair pathway. The two missing entries indicate that we could not find primary publications directly addressing the role of Nej1 in HR or Dnl4 in SSA, but neither would be considered to be likely. BIR, break-induced repair; GC, gene conversion.

|  | NHEJ | HR | SSA | MMEJ |
| --- | --- | --- | --- | --- |
| Dnl4 | +<sup>36,37</sup> | -<sup>38</sup> |  | x<sup>39</sup> |
| Nej1 | +<sup>40</sup> |  | x<sup>41</sup> | x<sup>32</sup> |
| Rad1 | -<sup>42</sup> | -<sup>43</sup> | +<sup>44</sup> | +<sup>39</sup> |
| Pol32 | -<sup>45</sup> | + (BIR), - (GC)<sup>46,47</sup> | -<sup>48</sup> | x<sup>32</sup> |
| Rad51 | -<sup>42</sup> | +<sup>49</sup> | -<sup>50,51</sup> | -<sup>32</sup> |
| Rad52 | -<sup>42</sup> | +<sup>52</sup> | +<sup>53,54</sup> | -<sup>39</sup> |

Here, we present the results of three key tests to elucidate the biological function of DDC override in budding yeast:

1. We finely tune DDC override timing using an optogenetic expression system for *CDC5* in an override-incompetent (*cdc5-ad*) background to demonstrate that checkpoint override is beneficial for survival and that, more specifically, the wild-type override time is optimal.
2. By tracking the fate of the acentric DNA fragment using a different optogenetic expression system, we find that the statistical benefit hypothesis is implausible.
3. Using dynamic shut-down of different DNA break repair pathways at the time of override (including of all known pathways together), we show that a pathway satisfying the genetic requirements for MMEJ is responsible for making DDC override beneficial for cells.

Our study is the first in-depth investigation of the biological function of DDC override, avoiding ambiguities inherent in using constitutive genetic perturbations. We expect the conceptual and methodological tools that we develop to serve as a model for elucidating override in other checkpoints whose function, if any, is currently poorly understood.

## Results

### Experimental system

To control cell cycle progression, induce DSBs, and monitor repair, we genetically modified budding yeast, starting from previously created strains^20^: We had deleted *CLN1-3* in the W303 genetic background (with *RAD5* ^55^ corrected) and inserted a *MET3pr-CLN2* construct. These genetic modifications allow controlling cell cycle Start by adding or removing methionine (+/-M), respectively, in the cell culture medium. We used the *cln1-3*Δ *MET3pr-CLN2* system to synchronize haploid cells in G1 in order to ensure only one copy of each chromosome was present when we induced a DSB (described below). In these cells, efficient repair by homologous recombination is naturally hindered by the absence of a homologous chromosome. We further ensured that our genetic modifications did not unintentionally enable homologous recombination or bias DNA repair by avoiding duplicate genetic sequences with optimal plasmid design^56^ for pop-in-pop-out genome manipulation, which eliminated extraneous DNA, including selection markers.

We had further introduced a *GAL1pr-HO* construct in these strains, which allows controlling Ho nuclease levels based on the carbon source: induction in galactose (G), lack of induction in raffinose (R), or repression in glucose (D). Ho cuts its recognition sites with >90% efficiency within 1 hr^57,58^, and is rapidly degraded (10 min half-life)^59^ when no longer produced. The natural Ho cut site in the *MATα* locus was blunted by multiple silent mutations in the alpha1 gene (*MATα-syn*). In our strains, Ho could instead only cut an artificially inserted Ho recognition site, placed at different loci, e.g., *URA3*, depending on the experiment. We extensively used a cut site whose break we could detect by fluorescence: An Ho restriction site was fused to the coding sequence of a destabilized yellow fluorescent protein gene, yEVenus-PEST, and the construct was inserted between the promoter and coding sequence of the strongly expressed *ADH1* gene. This DSB sensor allowed us to induce one DSB only during a brief time window in G1, and then identify the cells that still had the DSB at a later time point by their low yellow fluorescence. (In flow cytometry, cell size was additionally drawn upon for discriminating between the two populations.) We did not need to enrich for cells with a DSB by disrupting wild-type DNA break repair or recutting the Ho cut site by continuous induction of *HO*. The strains used in this work thus had the basic genotype *cln1*Δ*0 cln2*Δ*0::MET3pr-CLN2 (promoter replacement) cln3*Δ*0::GAL1pr-HO HTB2-mCherry::HIS5 MATα-syn RAD5*, which we refer to as ET44. The strains further either had an Ho cut site at *URA3*, in the DSB sensor at *ADH1* (*ADH1pr-HOcs-yEVenus-PEST-ADH1*), or in a similar mCherry-based sensor at *TDH3* (*TDH3pr-HOcs-mCherry-PEST-TDH3*). Further genetic modifications are indicated.

The basic protocol for our experiments^20^ encompassed overnight growth in synthetic complete (SC) R-M medium, 2-hour arrest in R+M to collect cells in G1, and 1-hour *GAL1pr-HO* induction in RG+M medium in order to create a DSB. At this point, in experiments in which DNA repair was permitted, we induced cell cycle Start by switching to D-M medium, which repressed *GAL1pr-HO*. When continuous recutting of the Ho recognition site was desired, we instead only initiated cell cycle start in RG-M medium. Approximately one hour later, cells budded reliably and arrested at the DDC, i.e., before anaphase^20^. When using DSB sensors, fluorescence increased markedly if the break was repaired, and stayed low otherwise, including during DDC override^20^. Both by fluorescence microscopy and flow cytometry, the two populations – with or without a DSB – could be easily distinguished. We deviated from our previous methodology^20^, in only sorting for cells with a DSB by fluorescence-activated cell sorting (FACS) once – not twice. We had sorted twice previously in order reduce the number of cells with uncut chromosomes but low fluorescence, which might erroneously have been selected as arrested with a DSB at the DDC. The drawback of double sorting is that it substantially reduced throughput as well as overall survival probabilities. Instead, here, we only sorted once in FACS experiments and used appropriate controls to draw conclusions based on relative changes.

### Wild-type override timing is optimal for cellular fitness

To demonstrate the functional relevance of DDC override, we wished to avoid confounding factors that arise when comparing genetically different strains, e.g., override-competent *CDC5* versus and -incompetent *cdc5-ad* cells. Furthermore, we wished to keep the genetic background the same in our comparisons and avoid DNA repair mutations. Instead, we opted for tuning the override time, quantifying survival probabilities, and identifying a potentially optimal override time (**Fig. 1**). We introduced the *cdc5-ad* mutation into our basic strain (ET44) as well as a system for optogenetic induction of *CDC5*, which allowed triggering override^60^ at specific times (*LIP-CDC5 PGK1pr-EL222*). The *cdc5-ad* mutation has not been observed to influence viability, cell cycle timing, or early post-break DSB repairs^20,21,26^. In wild-type cells, *CDC5* expression increases during DDC arrest, which is essential for DDC override^60,61^. Thus, induced *CDC5* expression mimics a critical step in the natural DDC override process. To control *CDC5* on demand, optogenetic expression was particularly well suited as it affords high temporal control without affecting metabolic pathways, in contrast to nutrient-controlled promoters such as the *MET3* or *GAL* promoters. We further chose the blue-light controlled LOV transcription factor El222^62^, which is comparatively tight and has a maximal strength of approximately 0.75 maxGAL1^63,64^.

**Figure 1:**
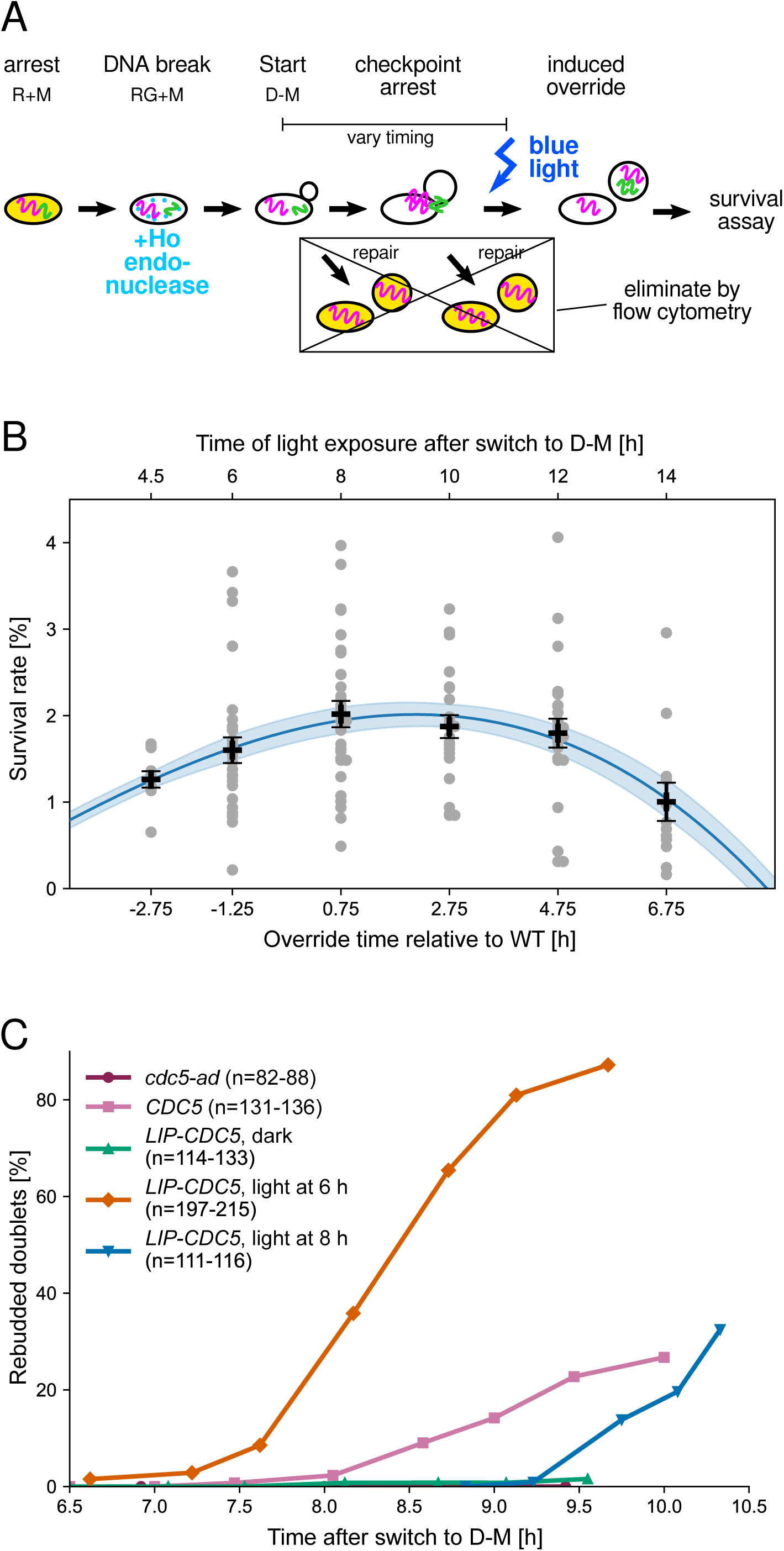
Tuning the DDC override time reveals the optimal override time for survival to be close to the wild-type override time. **A.** Protocol using ET44 cdc5-ad LIP-CDC5 PGK1pr-EL222 ADH1pr-HOcs-yEVenus-PEST-ADH1 cells. Starting with a log-phase culture in R-M medium, cells were collected in a G1 arrest in R+M for 2 hrs. GAL1pr-HO was induced for 1 hr in RG+M medium, and then the cell cycle was restarted and *GAL1pr-HO* was suppressed by switching to D-M. After 5 hrs (or 4 hrs for the first light induction point, see main text), large and low-fluorescence cells were selected by FACS and spread on D-M agar plates. A light pulse was administered at the time points indicated on the top horizontal axis in panel B. **B**. Survival rate versus induced override time after switching to D-M (top horizontal axis) or relative to the wild-type over-ride time (bottom horizontal axis), see panel C. Each grey point represents the fraction of colony-forming cells on one plate. Black points and error bars represent the mean and the standard error of the mean (SEM), respectively. The blue line is a cubic spline fit through the mean, and the shaded region represents the interpolated SEM from the fit. 8 repetitions were combined with interday normalization (**Methods**). **C**. On plates, the time to rebudding of mother-daughter doublets was scored as a function of time by brightfield microscopy. By comparing to wild-type override timing with *CDC5* (with no light-controlled *CDC5*), we related light-induced override to the wild-type override time by comparing when 15% of arrested mother-daughter doublets rebudded.

We extended our basic protocol by using FACS to select large and low-fluorescence cells at 5 hrs (4 hrs for the first time point, **Methods**) after the switch to D-M (**Fig. 1** A). The cells were spread onto D-M plates and were exposed to blue light at different time points (top horizontal axis, **Fig. 1** B). After 2-3 days, we counted the fraction of plated cells that formed colonies, which differed depending on the time of *CDC5* induction; survival peaked when *CDC5* was induced at 8 hrs after release into D-M. To calibrate the light-induced override times with respect to the wild-type override time, we compared the timing of mother and daughter rebudding after plating (**Fig. 1** C). Translating the induced override times to the natural override time (with unperturbed, wild-type *CDC5*), we observed the optimum override time was very close to the wild-time override time (peak survival is at +0.75 hrs with respect to the wild-type override time).

### Chromosome fragment segregation pattern conflicts with statistical benefit hypothesis

To elucidate the mechanism rendering override beneficial, we began by investigating the statistical benefit hypothesis. For replication and segregation of the genetic material into different cells to increase the overall chance of repair, the fate of the broken chromosome is critical. Specifically, the centromeric and acentric (i.e., centromere-less) chromosome fragments must end up in the same cell for repair to be possible. If a cell does not inherit the acentric fragment, its lineage will generally be dead, at a minimum, due to the loss of essential genes (essential regions predominate over deletable regions in the budding yeast genome^65^) and of the centric fragment’s telomere, which has a low chance of being regenerated. If the acentric fragment does not segregate into both mother and daughter cells after override, the genetic material would not be multiplied in a manner that would increase the rate of repair purely probabilistically.

To track the fate of the acentric fragment, we modified our basic strain genotype and protocol in the following ways (**Fig. 2** A): We induced *GAL1pr-HO* in galactose medium for a longer period (5 hrs) before switching to D-M medium. During this time, Ho repeatedly cuts its recognition site except if the chromosome fragments are rejoined by error-prone repair, which is too rare to occur in the relatively small number of cells followed here longitudinally by timelapse microscopy. After long *HO* induction, the cells turned out to no longer repair the DSB after *HO* was re-repressed, allowing us to track broken chromosome fragments efficiently in glucose medium; cells remained arrested at the DDC and overrode the checkpoint with typical wild-type timing of 8.2±0.5 h (mean±SEM, **Fig. 2** B). (A similar analysis with our previous arrest times and release into G-M yielded similar results, not shown.)

**Figure 2:**
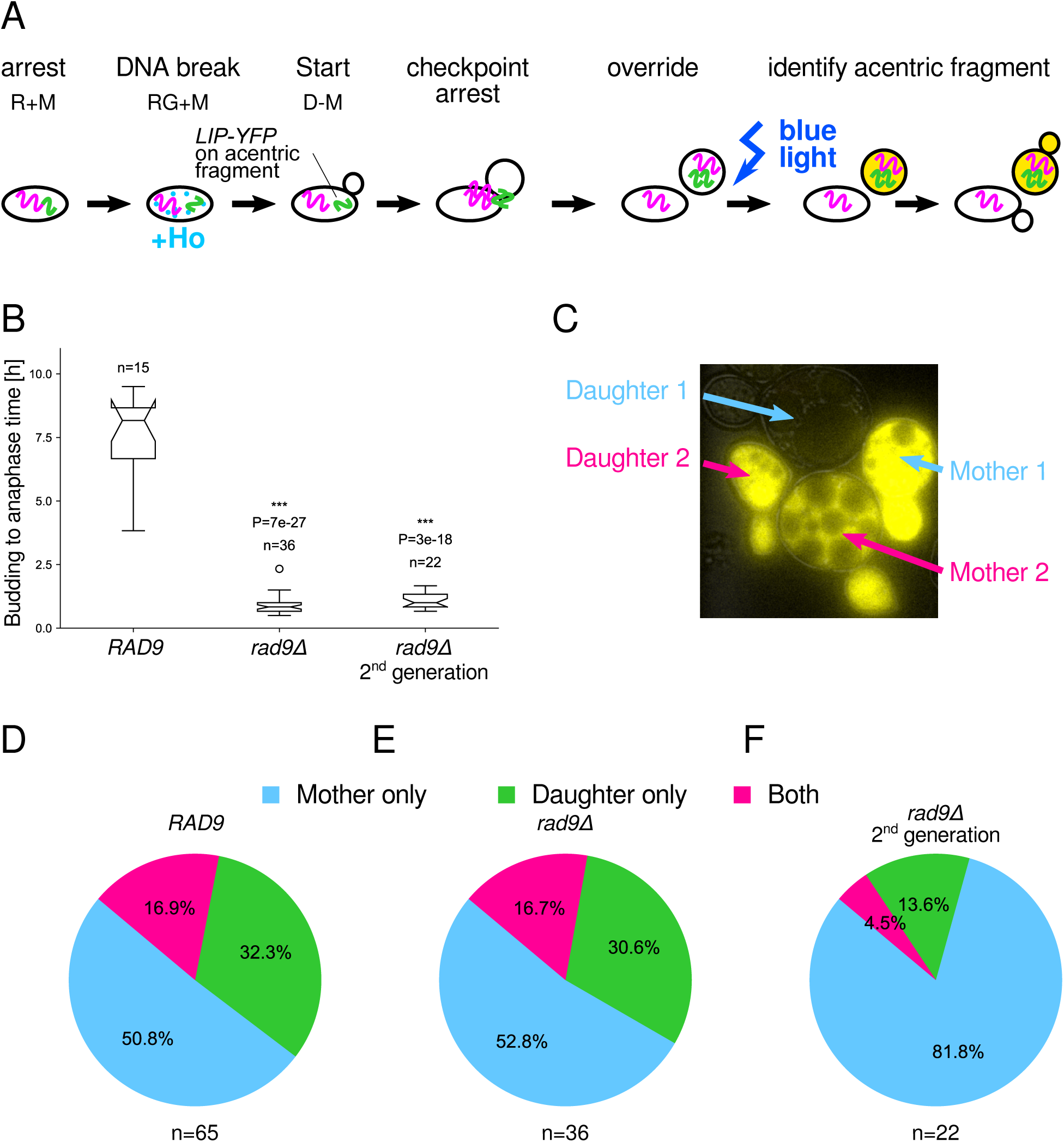
The acentric chromosome fragments co-segregate. **A**. Experimental protocol. ET44 *ura3*Δ*::HOcs::KanMX* cells were arrested in R+M for 2 hrs, as before, and switched to RG+M for 5 hrs. Once released in D-M medium, cells budded, arrested pre-anaphase for extended period (panel B), and then overrode the checkpoint. At 9 hrs after release into D-M medium, these cells, which also carried *PGK1::PGK1pr-EL222* and a *LIP-yEVenus* construct centromere-distal to the Ho cut site, were exposed to blue light. The mother and daughter cell that inherited the acentric fragment were identified by expression of the optogenetic yEVenus reporter. **B**. DDC override statistics in experiments presented in this figure. **C**. Two (of three) outcomes after override, where the ‘mother 1’ cell kept the acentric fragments (top, blue) or both ‘mother 2’ and ‘daughter 2’ cells have the acentric fragment (bottom, magenta). **D-F**. Inheritance patterns after the first override with a wild-type *RAD9* gene (D), after the first override with a *rad9*Δ deletion (E), or after an additional mitosis with a *rad9*Δ deletion (F).

We used an Ho recognition site at *URA3* on chromosome V instead of our fluorescent DSB reporters because we were not primarily interested in DSB repair, which did not occur here. Instead, we inserted the yellow fluorescent reporter under the control of the light-inducible promoter (LIP) controlled by El222 at a distance of 91.5 kbp telomere-proximal from *URA3*. We exposed cells to blue light at 9 hrs after switching to D-M medium, which allowed us to detect by light-induced optogenetic expression of yEVenus whether mother, daughter, or both had the acentric fragment (**Fig. 2** C). We found that in only 16.9% of cases both mother and daughter had the acentric fragment (**Fig. 2** D). In the other 83.1% of cases only one of the progeny had the acentric fragments, with retention in the mother cell being more prevalent. The low likelihood of independent segregation events argues for the replicated acentric DNA fragments segregating together. Our co-segregation pattern agrees qualitatively with previous observations^66^, which, however, used a GFP-LacI/LacO system to track chromosome fragments, whose fluorescent puncta are challenging to detect, and scored cells in a high-Clb2dbΔ mitotic exit arrest, which could influence segregation. A mother-cell bias was not observed in this work^66^ but has previously been seen in acentric ARS plasmids^67^.

Next, we wished to examine how the acentric fragments segregate in the next cell cycle since the association of the acentric fragments may be regulated, e.g., by *RAD52* ^66^. Therefore, the next cell cycles could segregate the acentric fragments differently during the first override. To assess this possibility, we wished to track the acentric fragment through the next anaphase. However, the very long and variable arrests before the first override (**Fig. 2** B) were followed by another slow cell cycle with highly variable timing (not shown). This overall high level of variability made it difficult to repeat the same experimental steps as before and merely delay activation of the optogenetic reporter. Instead, we shortened the first checkpoint arrest using the *rad9*Δ deletion. While the first post-break budding-to-anaphase time was much shorter now, 0.83±0.05 h (**Fig. 2** B), the acentric fragment segregated very similarly as in *RAD9* cells (**Fig. 2** E). In the subsequent cell cycle, which followed immediately and was also rapid (1.0±0.1 h, **Fig. 2** B), we observed again that the vast majority of acentric fragments did not segregate independently (**Fig. 2** F). Overall, these data make it implausible that replication and segregation of the broken chromosome lead to increased chances for repair and survival purely probabilistically.

### Dynamic perturbations reveal repair pathways increasing override-associated fitness

Next, we explored the repair benefit hypothesis. We knocked down specific DNA repair proteins in a temporally precise manner – after DDC arrest but before override – in order to evaluate their potential contribution to the override benefit. We did not use constitutive mutations in DNA repair genes in order to avoid any possible interference in DSB processing, early repairs, preparatory steps needed for repairs, or the proper establishment of DDC arrest. Instead, we selected key genes in DSB repair pathways (**Table 1**) and tagged them with the AID* sequence^68^ for on-demand knock-down. The AID* peptide is a truncated version of the auxin-inducible degron (AID) and guides Tir1 to target AID*-tagged proteins for degradation in the presence of auxin or 1-Naphthaleneacetic acid (NAA). To reduce leakiness in the absence of ligand, we further placed *TIR1* under the control of the rtTA/*tetOpr* system, benchmarked previously^63^.

We used the protocol illustrated in **Fig. 3** A: We arrested cells proliferating in R-M medium by switching to R+M for 2 hrs, expressed *GAL1pr-HO* for one hour in RG+M medium, and then released cells from cell cycle arrest and re-repressed *GAL1pr-HO* in D-M medium. After 2 hrs, we added doxycycline (dox) and NAA to the cell culture in order to degrade AID*-tagged repair proteins – post arrest but pre override. (Control cells were not exposed to doxycycline and NAA.)

**Figure 3:**
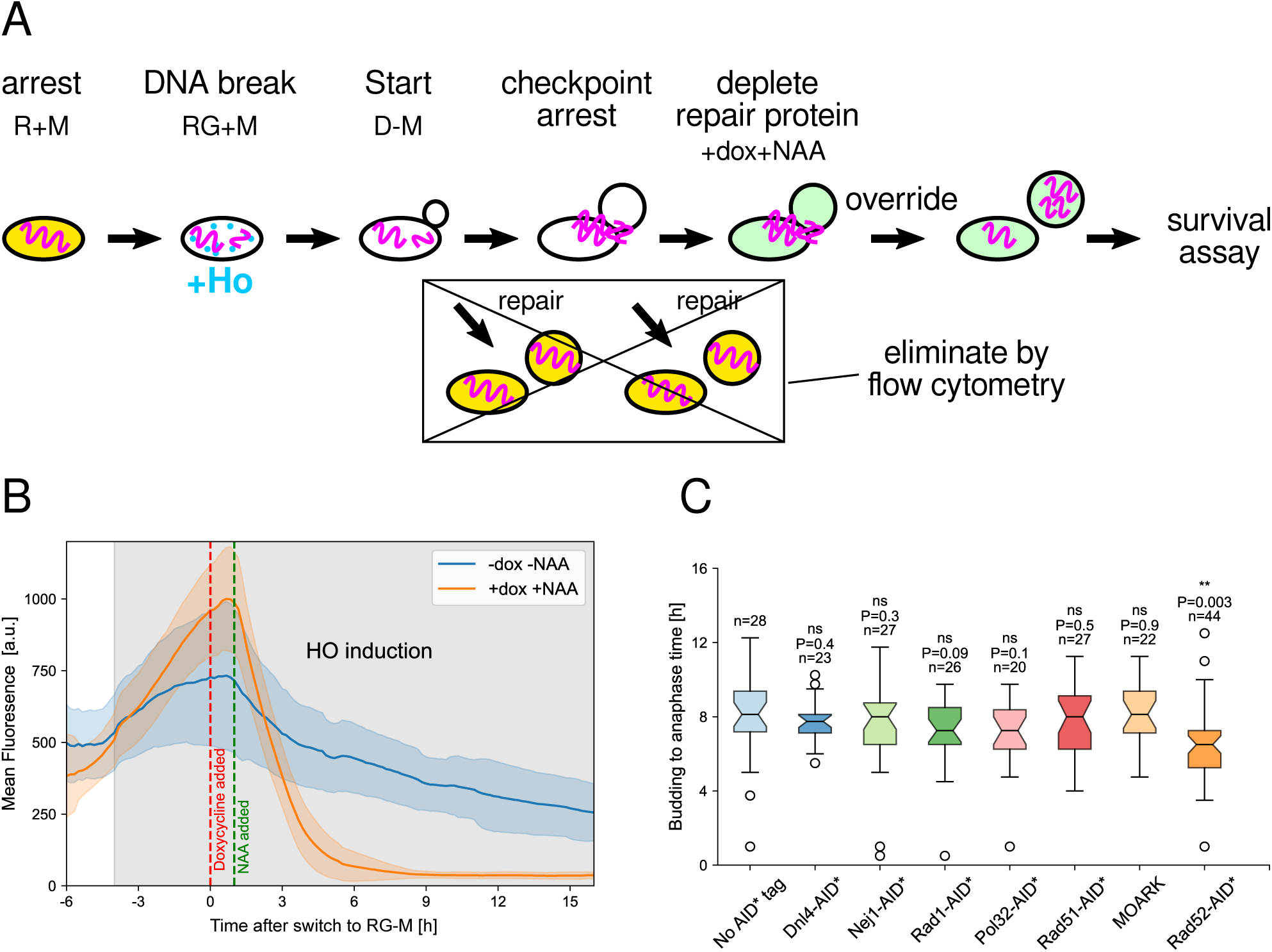
Dynamic repair pathway knockdown using Tir1/AID*. **A**. ET44 cells with the yEVenus DSB reporter at *ADH1* were arrested in R+M medium for 2 hrs, *GAL1pr-HO* was induced for 1 hr in RG+M medium, and the cell cycle was re-started by switching to D-M medium. At 2 hrs after switching to D-M, doxycycline and NAA were added in order to degrade AID*-tagged repair proteins. At 5 hrs, FACS was used to sort for large and low-fluorescence cells, which were plated on D-M+dox+NAA agar plates. After 2-3 days, the fraction of colony-forming cells were counted. **B**. Quantification of the efficiency of Tir1/AID* in our system. ET44 *TDH3pr-HOcs-mCherry-PEST-TDH3 TetOpr-SKP1-TIR1 ADH1pr-AID*-yeGFP* cells, proliferating in log phase in R-M medium, were arrested in R+M for 2 hrs at the beginning of the fluorescence timelapse recording, *GAL1pr-HO* was induced for 4 hrs in RG+M medium, and the cell cycle was restarted while continuing to express *HO* continuously in RG-M medium. To test the efficiency of the Tir1/AID* degradation system, doxycycline and NAA were added to the medium as indicated (orange trace, mean of n=29 cells) but not for control cells (blue trace, mean of n=18 cells). Orange and blue shading indicate the standard deviation (STD). **C**. Repair proteins knock-downs have no or a small effect on override times (budding to anaphase) measured in microfluidic chips by fluorescence microscopy. The same protocol was followed as for the bulk cell culture experiments except cell cycle Start was initiated by switching to RG-M medium instead of D-M to maintain a DSB in all cells.

Using the cells’ yEVenus DSB sensor at *ADH1* and FACS sorting, cells with a DSB were selected 5 hrs after switching to D-M and spread on D-M+dox+NAA agar plates (controls on D-M plates without doxycycline and NAA). The fraction of colony-forming cells was counted after 2-3 days.

We verified that the timing of inducible degradation was effective for the repair pathway knock-down experiments. Using our basic strain genotype with a cut site at *TDH3* and an AID*-tagged yeGFP reporter, we found that doxycycline-induced transcription of *TIR1* followed by NAA-induced activation led to 90% reduction of the yeGFP signal within 4 hrs during the DSB arrest (**Fig. 3** B). Thus, adding doxycycline and NAA at 2 hrs after release from cell cycle arrest in D-M in our repair knock-down experiments should reduce the AID*-tagged repair proteins by over 90% right before cells start to override (beginning approx. 6 hrs after release in D-M). We further verified that the repair pathway knock-downs did not affect override (**Fig. 3** C), being mildly sped up with Rad52-AID* knock-down.

To assess the effect of each repair pathway knock-down quantitatively, we used override-incompetent *cdc5-ad* cells as controls; an override-associated benefit requires the repair pathway knock-down to reduce the survival probability in override-competent (*CDC5*) cells more than in override-incompetent (*cdc5-ad*) cells. Otherwise, the repair protein is increasing the chance of late DSB repairs generally, independently of override.

We excluded that doxycycline-induced *TIR1* expression and activation by NAA affected survival if no protein was tagged with AID* (**Fig. 4** A). The observed difference between *CDC5* and *cdc5-ad* survival in this and the following experiments also supports the conclusion that DDC override is beneficial for survival, which we established in a more careful manner above (**Fig. 1**).

**Figure 4:**
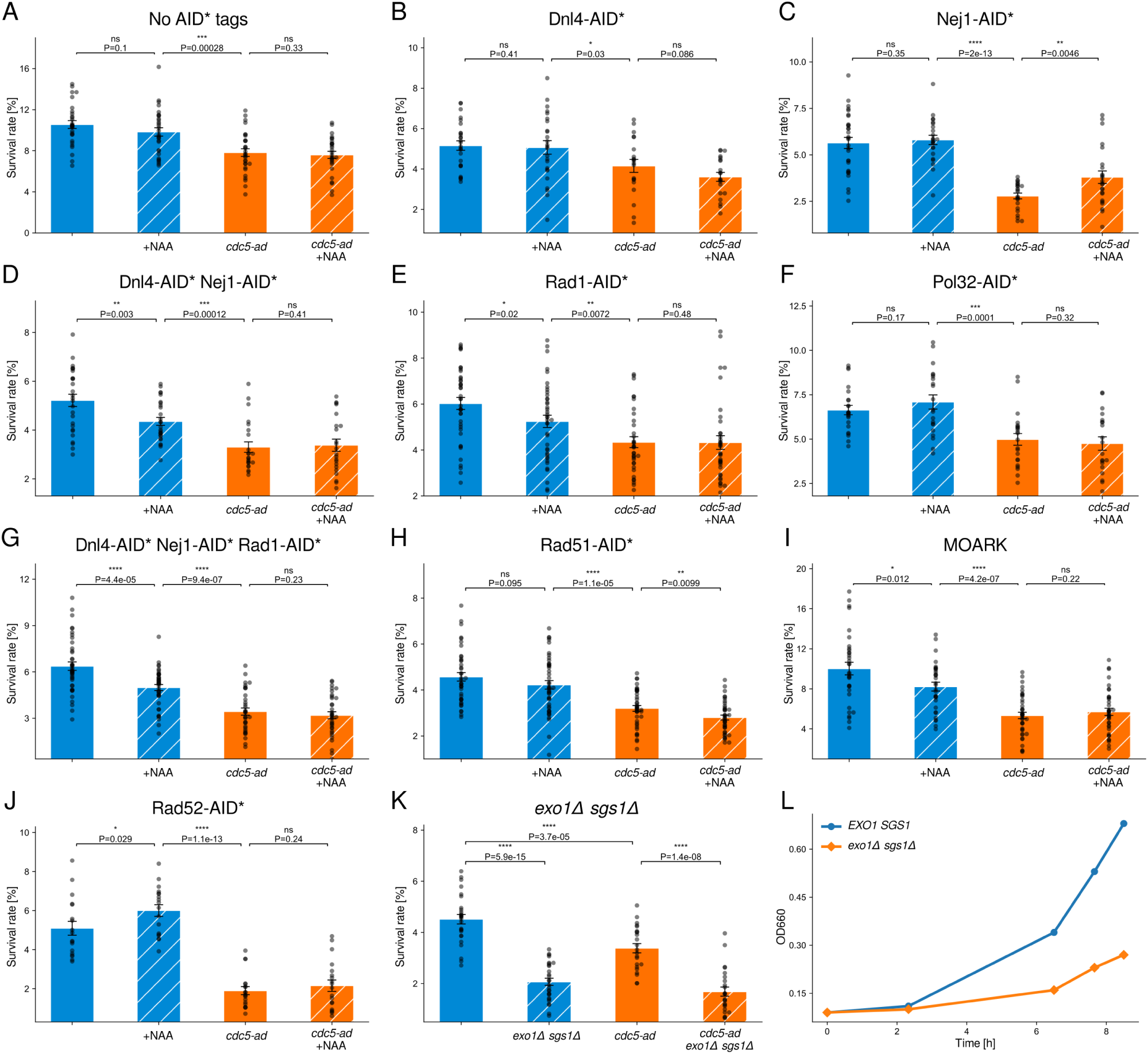
The repair pathway conferring a survival benefit to DDC override revealed by dynamic knock-down experiments. **A-J.** Fraction of surviving, colony-forming cells with AID*-tagged proteins, as indicated. +NAA indicates that doxycycline and NAA were added 2 hrs after cell cycle release in D-M. **K-L**. Fraction of surviving, colony-forming cells (K) or optical density in D-M liquid culture (L) with constitutive *exo1*Δ *sgs1*Δ deletions. Each circle represents the result of one measurement, that is, one agar plate. The height of the bars indicates the mean and error bars represent the standard error of the mean (SEM).

First, we tested whether c-NHEJ could provide an override-associated survival benefit. When we knocked down Dnl4-AID* or Nej1-AID* individually starting at 2 hrs after the switch to D-M, we saw no significant difference compared to the survival rate without knockdown (**Fig. 4** B, C). However, when we combined the two knockdowns, we saw a significant decrease in survival (**Fig. 4** D), accounting for 45±5% (mean±SEM) of the difference in survival between *CDC5* and *cdc5-ad* cells. We deduced that either c-NHEJ is responsible for a survival benefit but two incomplete knock-downs of Dnl4-AID* and Nej1-AID* were needed to shut this repair pathway down sufficiently to affect survival, or the knock-downs were effective but the pathway is not fully dependent on either protein individually, e.g., as in MMEJ (**Table 1**). To test directly whether a pathway with the requirements of MMEJ was needed for the survival benefit, we knocked down Rad1-AID* post establishment of the DDC but pre override. We observed a similar decrease in survival (**Fig. 4** E), accounting for 46±4% (mean±SEM) of the difference of survival between *CDC5* and *cdc5-ad* cells. To further probe the case for MMEJ, we knocked down Pol32-AID* (**Fig. 4** F). This knock-down had no significant effect on survival, which can either be due to sufficient Pol32 molecules remaining to support MMEJ, no DNA synthesis being required for this particular repair product, or the affected pathway being actually independent of Pol32.

The repair proteins that increased override-associated survival, Rad1 or combined Dnl4-Nej1, are usually associated with different pathways, MMEJ/SSA or c-NHEJ, respectively. To clarify whether two pathways contributed independently or only one pathway, we induced degradation of all three proteins together (**Fig. 4** G). Again, the survival benefit of overriding cells, now with Dnl4-AID*, Nej1-AID*, and Rad1-AID* knocked down simultaneously, showed a similar decrease as before, accounting for 47±5% (mean±SEM) of the difference of survival between *CDC5* and *cdc5-ad* cells. Given that about half of the benefit of override can be explained by knocking down Dnl4-AID* and Nej1-AID* combined, Rad1-AID* alone, or all three together, a repair pathway with the genetic requirements of MMEJ appears to be generating the fitness benefit of DDC override.

While HR is unlikely to play a role in increasing survival of haploid cells with a DSB, we nevertheless probed this possibility. As expected, we found no significant decrease in survival with a Rad51-AID* knock-down (**Fig. 4** H). To test the relationships of all the pathways affected by the repair proteins discussed thus far, we tagged all five proteins (Dnl4, Nej1, Pol32, Rad1, Rad51) with AID* in the ‘mother of all repair knock-down’ (MOARK) strain (**Fig. 4** I). The MOARK strain showed a similar override-associated survival benefit, 38±5% (mean±SEM), as already observed with the triple knock-down suggesting no further synergy, compensation, or back-up functionality.

Finally, we sought to further investigate whether the repair pathway increasing override-associated survival was genetically more similar to MMEJ or, actually, SSA. Both processes are Rad1 dependent, which we observed in override-associated repair (**Fig. 4** E). However, a role for Dnl4 (**Fig. 4** D) has only been described, to our knowledge, for MMEJ^39^, and a Rad51 knock-down should have benefitted SSA^51^ but only a statistically non-significant decrease in survival was observed (**Fig. 4** H). On the other hand, the lack of Pol32 dependence raises questions regarding MMEJ, although multiple explanations for this observation can be given, including residual Pol32 activity. Thus, we focused on a further distinction between SSA and MMEJ, the requirement for *RAD52*. Rad52 has been suggested to have a role in recognizing the presence of unrepaired DNA and thus may be implicated in the decision to override^69^. This may explain why cells with a Rad52 knockdown override faster than wild type (**Fig. 3** C). The difference in override timing when Rad52 was degraded was the main reason that we did not also tag *RAD52* with *AID\** in the MOARK strain; a reduction in survival in the Rad52-AID* knock-down would be inconclusive since it could be due to the impairment of Rad52-dependent repair pathways or, alternatively, to premature override. To test the effect of Rad52 degradation strictly to distinguish MMEJ from SSA, we used a previously validated AID*-tagging strategy^70^ and knocked down Rad52-AID* dynamically as well (**Fig. 4** J). We observed a higher survival rate in *CDC5* cells, which bypasses a potential early-override survival penalty and is in line with MMEJ but not SSA. The increased MMEJ efficiency with Rad52-AID* knockdown is in agreement with previous results with a *rad52*Δ deletion^71^.

Exploring the mechanism conferring a benefit to DDC override further faces challenges due to the exacting nature of our experiments. Testing whether long-range resection affected the override-associated survival benefit, we observed a substantial decrease in survival in both *CDC5* and *cdc5- ad* cells with *exo1*Δ *sgs1*Δ double deletions (**Fig. 4** K). However, the decline was not specific to override-specific repairs or late repairs since the *exo1*Δ *sgs1*Δ cells proliferated overall clearly poorly (**Fig. 4** L).

Interpreting the repair process based on the sequences of repaired DSB loci in surviving colonies also faces challenges inherent to our experimental requirements. The cut sites in 57 colonies emerging from surviving cells across the different experiments (26 colonies from -dox-NAA and 31 from +dox+NAA conditions in **Fig. 4**) showed no modifications to the Ho recognition site sequence (TTCAGCTTTCCGCAACAGTATAATTTTATA) or the wider locus. Error-free repair is in apparent contradiction with the perceived association of MMEJ with imprecise repairs, particularly deletions^39,71,72^. However, in past work, cut sites were generally analyzed in colonies formed from a founder cell that survived continuous endonuclease expression, which foreclosed error-free repair^39,71,72^. We implemented multiple measures, including a fluorescent DSB sensor and a multi-step experimental protocol involving FACS, precisely to avoid continuous recutting and examine unimpeded repair processes.

Together, our results show that DDC override gives cells a fitness advantage due to repair processes involving MMEJ factors. A new experimental methodology combining a DSB sensor, FACS, and repair pathway knock-downs were needed to observe these effects.

## Discussion

Checkpoint override unblocks biological processes in the presence of persistent damage, yet the functional consequences of this fundamental phenomenon have not been established; quantitative approaches are needed based on careful perturbations and measurements. In our study, we answered this question in budding yeast cells. By finely tuning the override time, we showed that the wild-type override time was optimal for survival (**Fig. 1**), avoiding comparisons of different genetic backgrounds. The simplest mechanistic explanation, the Statistical Benefit Hypothesis, was rendered implausible by the segregation pattern of the acentric chromosome fragments, which we tracked using an optogenetic marker and which we found travel together in the first and second post-arrest mitosis – the latter demonstrated in a *rad9*Δ background (**Fig. 2**). Note that several modes of acentric fragment segregation exist in other organisms^73^, raising the possibility that increasing the chance of repair purely probabilistically is relevant in other contexts. Instead, dynamic knock-down experiments showed that override-associated repair confers an increased chance of survival, supporting the Repair Benefit Hypothesis (**Figs. 3, 4**).

The relationship between survival rate and override timing (**Fig. 1**) likely represents a more intricate interplay of factors than may appear at first glance. Both earlier and later override than in wild type are detrimental to survival. Regarding the decrease of survival rate with later override, we speculate that cell cycle-arrested cells invariably lose viability, which is seen in different contexts^74^. On the other hand, understanding why earlier override lowers the chance of survival likely requires precise and systematic quantification of the kinetics of different DNA repair pathways. In previous work, we measured the chance of pre-override DSB repair kinetics, which declines with time^20^. In addition, we quantified the chance of surviving DDC override and modeled it as constant in time. So, as a cell with DNA damage remains arrested, pre-override repair decreases in likelihood and override becomes eventually more beneficial. Combined with this prior analysis, we propose that the efficiency of other repair pathways, e.g., c-NHEJ, declines more rapidly than override-associated repair requiring MMEJ factors. Interestingly, different repair pathways and mechanisms have been seen to be particularly effective within specific, multi-hour-long time windows: BIR takes more than ≈6 hrs after a DSB due to slow initiation of new DNA synthesis^47^ and Rad51 filament completes about 90% of the search for a homologous sequences within ≈6-8 hrs^75^. More relevant to our study, MMEJ repair was found to be very slow, peaking 6-8 hrs after DSB induction, which is thought to be due to the instability of microhomology annealing^76^. Thus, the peak in survival rate (**Fig. 1**) conforms with previously established DNA repair timescales.

Given that the mechanisms of DDC override are not fully understood themselves, the molecular intersection of override with DSB repair can only be conjectured. One potential focal point is *SAE2*, which is necessary for DDC override as measured by microcolony formation in the presence of one DSB and whose overexpression suppresses DDC arrest^77,78^. Sae2 is further phosphorylated by Cdc5^61^ and is needed for SSA^79^ and MMEJ^69^. Relatedly, in mammalian cells, at least two mechanisms link MMEJ to mitosis and PLK1 (Cdc5 in yeast): PLK1 phosphorylates CtIP (Sae2 in yeast) and PolΘ (no yeast homolog) to promote MMEJ^80,81^. While a model by which prolonged arrest and PLK1/Cdc5 activity lead to increased MMEJ activity is appealing, the role of override would be entirely unclear.

Despite long-standing interest in override of checkpoints generally, and of the DDC checkpoint in budding yeast in particular, its overall functional relevance had remained unclear. While there were conceptual and technical challenges to solving this question, e.g., a methodology that does not block repair but merely selects cells that maintain a DSB for many hours, the size of the effect of a few percent (**Fig. 4**) may also have contributed to the challenge of solving this problem. However, from the point of evolutionary selection, a survival advantage of a few percent suffices, considering the large size of yeast populations and their evolutionary age. For comparison, the spindle assembly checkpoint (SAC) corrects far fewer errors in budding yeast in the absence of spindle poisons^82^, yet, has clearly also evolved and been maintained.

While repairs substantially contribute to the benefit of overriding the DDC (**Fig. 4**), the gap in survival rate between *CDC5* and *cdc5-ad* cells is not fully explained. One reason may be residual repair protein activity, especially in cells that override relatively early. We timed doxycycline and NAA supplementation so that AID*-tagged proteins should be reduced, on average, by over 90% at the start of checkpoint override (**Fig. 3** B). However, the minimum number of protein molecules to effectuate repair is not known. Furthermore, degradation of AID*-yeGFP slowed down after 4 hrs (**Fig. 3** B), suggesting limitations to improving inducible degradation.

Our work introduces new concepts, tools, and results that should guide the interrogation of check-point override more broadly. In many systems such as proliferating unicellular organisms, germline cells, embryonic cells, and developmental processes, evolution can be expected to have shaped a trade-off between risk and speed^20^. Arresting at a checkpoint reduces reproductive potential, while override increases the risk of failure. In such contexts, optimally timed override should have adaptive value. Interestingly, fail-safe mechanisms have also been described for the cell cycle oscillator in bud-ding yeast^83,84^, reducing the chances of catastrophic failure. By contrast, the growing evidence for checkpoint override or “leakiness” in non-cancerous human somatic cells presents a puzzle. Given the risks that genomic instability poses to the organism as a whole, it remains unclear whether such override reflects evolutionary tolerance, molecular noise, or a presently unrecognized physiological benefit. The study of checkpoint override thus remains intellectually provocative, demanding innovative methods to illuminate a phenomenon at the intersection of fundamental molecular biology, evolution, and potentially disease.

## Methods

### Strains

All strains have the W303 strain background with *RAD5* corrected. Most genetic modifications were performed with the *URA3* -pop-in/5-FOA-pop-out method^56^ to reduce the introduction of extraneous DNA, or transformation with an integrating plasmid, as indicated. The standard LiAc method was used for budding yeast transformation. Plasmids were created with standard restriction cloning or ordered from Twist Biosciences. A full list of plasmids and strains used in this study can be found in the Supplementary Tables 1 and 2.

### Basic preparatory protocol for bulk culture experiments

Strains were inoculated in synthetic complete medium with 3% raffinose and without methionine (R-M) until they reached stationary phase. The day before the experiment, a small volume of cell culture was transferred to fresh medium in an amount calculated to reach early log phase the next morning. The next morning, samples were transferred to a flask and 100 mg/L (5x) methionine was added. The flasks were incubated for 2 hrs after which 3% galactose (G) was added. The flasks were incubated for another hour. The cells were spun down at 700 g for 3 min, the medium was decanted, cells were rinsed with synthetic complete medium with 2% glucose and without methionine (D-M), spun down at 700 g for 3 min, the medium was decanted again, and D-M medium was added. Depending on the experiment, 100 mg/L methionine was added after 1.5 hrs to prevent repaired cells from overcrowding and the flasks were incubated until the next steps. Incubations were done in a shaking incubator set to 250 rpm and 30 ^◦^C.

### Flow cytometry

At 5 hrs after the removal of galactose, samples were spun down to concentrate cells. We used a Sony SH800 instrument for FACS. 2500 cells were sorted into new tubes containing 100 *µ*L of D-M medium. Directly after sorting all samples, the tubes were quickly spun down to collect all droplets, plated on D-M agar plates, and incubated for 2-3 days. All plates were in the incubator by 6 hrs post removal of galactose at the latest.

### Light induction

After spreading cells on plates, they were exposed to 30 min of 30 sec on, 30 sec off pulsing blue light at different time points (4.5, 6, 8, 10, 12, and 14 hrs after switch to D-M) using commercial blue LED panels (HQRP, USA) placed 20 cm above the plates. For the 4.5 h time point, the samples were sorted by FACS at 4 hrs instead of 5 hrs after the switch to D-M medium and exposed to light right after plating. For normalization and comparison with the other data, some samples sorted at 4 hrs were exposed to light at 6 hrs. The 4 hr sorting results were then normalized so that the 6 hr light exposure result matched that with 5 hr sorting.

### Auxin-inducible degradation

At 2 hrs post removal of galactose, the samples were split into two equal parts in two different flasks. In one flask, 0.02 mM NAA and 1 *µ*M doxycycline were added. The samples were returned to the shaking incubator and FACS was performed as described above. The sorting tubes and the plates used for the +dox+NAA samples had the same concentration of NAA and doxycycline in/on them.

### Imaging and image processing

Timelapse images were recorded with a Plan Apo *λ* 60x/1.40 oil objective and a Hamamatsu Orca-Flash4.0 camera. Cells were grown in CellASIC microfluidic chips. Images have 16 bit depth. The diascopic light was generated by Nikon Ti2-E LEDs. The exposure time was 100 ms. Images were processed with YeaZ^85^.

### Data analysis

Survival rate measurements were performed with most time points (**Fig. 1**) or all conditions to be compared (**Fig. 4**) included in a set of experiments on a given day. To compensate for day-to-day variation, we normalized each day’s mean to the mean over all related experiments. Data points whose z-scores after normalization where more than 3 or less than -3 were removed. P-values were determined using a one-tailed t-test. Error bars denote the standard error of the mean (SEM), unless noted otherwise. For the time point experiments (**Fig. 1** B), the fit was found using a non-linear least squares fitting of a cubic polynomial. The error of the fit was linearly interpolated between the measured SEM values at consecutive data points.

## Acknowledgments

We thank Ahmad Sadeghi and Roxane Dervey for creating reagents, help performing experiments, and early data acquisition. We thank Profs. Susan Gasser, Galit Lahav, Timothy Mitchison, and Marcus Smolka for advice and fruitful discussions. Support for RO was provided by SNSF grants CRSK-3 190526, 310030 204938, and CRSK-3 221036 awarded to SJR.

## Author contributions

RO performed the experiments and analyzed the data. RO and SJR wrote the manuscript. SJR conceptualized the research, supervised the worked, and acquired funding.

